# Circadian rhythms regulate refractive development across species

**DOI:** 10.64898/2026.04.02.713440

**Authors:** Teele Palumaa, Nele Taba, Shruti Balamurugan, Angus C. Burns, Jakob M. Cherry, Shane P. D’Souza, Maris Teder-Laving, Estonian Biobank Research Team, Richa Saxena, Tõnu Esko, Erik Abner, Machelle T. Pardue, Priit Palta

**Affiliations:** Estonian Genome Centre, Institute of Genomics, University of Tartu, Tartu, Estonia; Department of Ophthalmology, Emory University, Atlanta, USA; Eye Clinic, East Tallinn Central Hospital, Tallinn, Estonia; Division of Sleep and Circadian Disorders, Brigham and Women’s Hospital, Boston, MA, USA; Program in Medical and Population Genetics, Broad Institute, Cambridge, MA, USA; Division of Sleep Medicine, Harvard Medical School, Boston, MA, USA; Center for Genomic Medicine, Massachusetts General Hospital, Boston, MA, USA; Division of Pediatric Ophthalmology, Cincinnati Children’s Hospital Medical Center, Cincinnati, OH, USA; Center for Visual and Neurocognitive Rehabilitation, Atlanta VA Health Care System, Decatur, GA, USA

## Abstract

Myopia is a rapidly escalating global public health challenge, yet the biological mechanisms linking modern lifestyles to abnormal eye growth remain unclear. Circadian rhythms have been implicated in refractive development, but causal evidence is limited. Here, we integrate population-scale human data with an experimental animal model to determine whether circadian misalignment contributes to myopia. In >265,000 individuals from the Estonian and UK Biobanks, late chronotype was consistently associated with myopia. To assess causality, we experimentally disrupted the alignment between behavioural and environmental rhythms in mice by housing them in non-24-hour light-dark schedules. Exposure to a lengthened cycle (T26) induced a myopic shift that was, notably, reversible in early adulthood. Retinal transcriptomics revealed enrichment of mitochondrial and hypoxia-related plasticity pathways, with transcriptional changes distributed across multiple retinal cell classes. Together, these findings identify circadian misalignment as a conserved and modifiable driver of myopia, highlighting opportunities for novel preventive and therapeutic approaches.

## Introduction

Myopia, or short-sightedness, is an ocular condition characterised by impaired distance vision. Its prevalence is currently estimated at 34-40% worldwide and is projected to affect 40-50% of the global population by 2050.^1,2^ Myopia usually manifests during childhood or early adulthood and, while correctable with optical methods, increases the risk of complications such as myopic macular degeneration, retinal detachment, glaucoma, and cataract,^3,4^ leading to substantial a societal and economic burden.^5,6^ Myopia results from disrupted refractive development, or emmetropisation, the process by which the growing eye coordinates axial length with the optical power of the cornea and lens to focus images sharply on the retina. Newborns are usually hyperopic, and emmetropisation gradually reduces hyperopia, ideally resulting in emmetropia, the refractive state in which parallel light rays are focused on the retina.^7–10^ Myopia arises when axial elongation exceeds this coordinated growth, causing images to focus in front of the retina.^8,11^ Hyperopia is often masked early in life by accommodation, but it often becomes symptomatic with age as lens elasticity declines.^9,11^

Accumulating evidence suggests that circadian rhythms influence refractive development. Genome-wide association studies (GWAS) have identified that genes associated with refractive errors are enriched for those involved in circadian rhythm regulation,^12^ and ocular parameters, such as axial length and choroidal thickness, exhibit prominent daily fluctuations.^13,14^ In animal models, experimentally induced myopia can disrupt or abolish these rhythms.^14,15^ Similarly, human studies suggest links between myopia and circadian rhythms, which have primarily been studied in the context of the sleep-wake cycle. Chronotype, or individual preferences for sleep and activity timing, range from early or morning types to late or evening types, and are shaped by several factors, including exposure to environmental stimuli, the intrinsic properties of the circadian clock, and its sensitivity to light.^16,17^ Studies examining sleep parameters, such as bedtime, sleep duration, and sleep quality, in relation to myopia have yielded inconsistent results (see reviews^18,19^). Some studies have reported a link between late chronotype and myopia,^20–22^ while others have found no associations.^23,24^ Notably, current studies are often limited by small sample sizes, inconsistent definitions of chronotype, and difficulty distinguishing correlation from causation.

Here, we addressed these gaps by first examining the relationship between chronotype and refractive error in a sample of over 265,000 individuals, leveraging two large cohorts: the Estonian and UK Biobanks. Motivated by these findings, we further investigated whether disrupting the alignment between the internal circadian clock and environmental timing cues can causally influence refractive development in mice.

## Results

### Association between chronotype and refractive errors in humans

We first examined the association between chronotype and refractive error in the Estonian Biobank. Chronotype was assessed using the Munich Chronotype Questionnaire (MCTQ), which defines chronotype as midsleep time on free days corrected for sleep debt (MSFsc).^16^ After rigorous quality control steps (**Supplementary Table 1**), 109,461 participants remained for analysis. Among them, 57.7% had no refractive error, 26.6% had myopia, and 15.7% had hyperopia (**Table 1**). As MSFsc showed a strong non-linear relationship with age (**Fig. 1a**), participants were grouped into age-specific MSFsc percentile categories (extreme early [P0-5], early [P5-25], intermediate [P25-75], late [P75-95], and extreme late [P95-100]), calculated within 1-year age bins. In an age- and sex-adjusted model (**Table 2**, **Model 1**), chronotype showed opposing associations with myopia and hyperopia: late chronotype was associated with higher odds of myopia and lower odds of hyperopia. The exact reverse pattern was observed for early chronotype. Given the strong link between education and refractive errors,^25,26^ we next fitted a model additionally adjusted for educational attainment. Although the corresponding effect sizes were modestly attenuated, the overall pattern remained robust (**Table 2**, **Model 2**, **Fig. 1b**). For myopia, extreme late chronotype was associated with increased odds of myopia (OR = 1.09, 95% CI: 1.02-1.16, *p* = 0.0085), whereas extreme early chronotype was associated with lower odds of myopia (OR = 0.91, 95% CI: 0.85-0.97, *p* = 0.0059). For hyperopia, the opposite was true: extreme early chronotype was linked with significantly higher odds of hyperopia (OR = 1.11, 95% CI: 1.02-1.20, *p* = 0.011), while the extreme late group had lower odds for hyperopia (OR = 0.88, 95% CI: 0.81-0.97, *p* = 0.0061). Intermediate late and intermediate early chronotype groups showed effects that lay between these extremes across all analyses.

**Figure 1.**
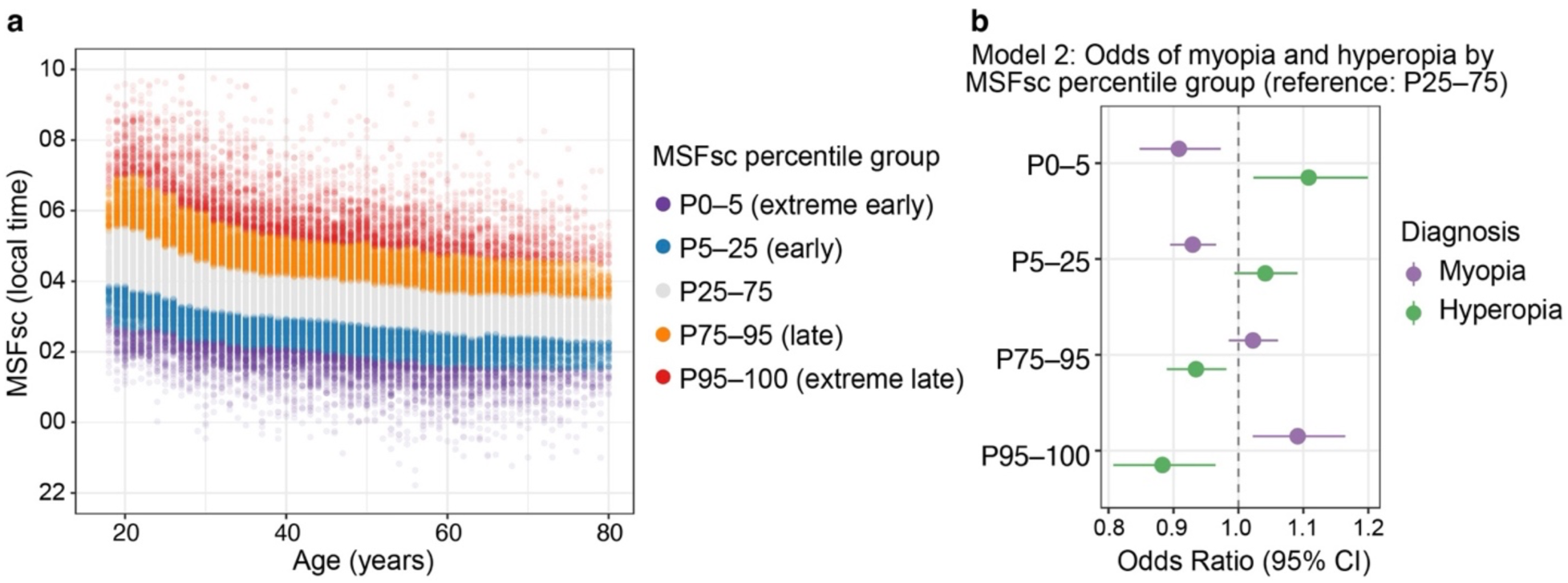
Chronotype distribution and associations with refractive error. (a) Distribution of midsleep on free days adjusted for sleep debt (MSFsc) by age in 109,461 participants from the Estonian Biobank. Percentile groups were defined within 1-year age bins: extreme early (P0-5), early (P5-25), middle chronotype group (P25-75), late (P75-95), and extreme late (P95-100). Each dot represents an individual, and the banding illustrates the age-specific distribution of chronotype across percentiles. (b) Odds ratios for myopia and hyperopia across MSFsc percentile groups in multivariable logistic regression (Model 2), adjusted for age, sex, and highest level of education. The P25-75 group served as the reference. Myopia risk was higher among late chronotypes, whereas hyperopia risk was higher among early chronotypes.

**Table 1.** Characteristics of the Estonian Biobank study population included in the logistic regression analysis.

| Characteristic | Overall<br>N = 109,461 (100%) | Refractive error |  |  | p value<sup>1</sup> |
| --- | --- | --- | --- | --- | --- |
|  |  | None<br>N = 63,113 (57.7%) | Myopia<br>N = 29,131 (26.6%) | Hyperopia<br>N = 17,217 (15.7%) |  |
| Age, median (Q1, Q3) | 43 (32, 56) | 42 (32, 53) | 39 (29, 51) | 57 (47, 66) | $<2 \times 10^{-308}$ |
| Sex, n (%) | | | | | $<2 \times 10^{-308}$ |
| Male | 36,453 (33.3%) | 24,408 (38.7%) | 7,224 (24.8%) | 4,821 (28.0%) |  |
| Female | 73,008 (66.7%) | 38,705 (61.3%) | 21,907 (75.2%) | 12,396 (72.0%) |  |
| Highest level of education, n (%) | | | | | $<2 \times 10^{-308}$ |
| Middle | 8,615 (7.9%) | 5,434 (8.6%) | 1,487 (5.1%) | 1,694 (9.9%) |  |
| Secondary | 50,072 (45.8%) | 29,122 (46.2%) | 11,743 (40.3%) | 9,207 (53.5%) |  |
| Higher | 50,664 (46.3%) | 28,493 (45.2%) | 15,875 (54.5%) | 6,296 (36.6%) |  |
| Unknown | 110 | 64 | 26 | 20 |  |
| MSFsc (hours from previous midnight), mean (SD) | 27.59 (1.14) | 27.61 (1.14) | 27.75 (1.16) | 27.26 (1.07) | $<2 \times 10^{-308}$ |

<sup>1</sup> To assess the differences in characteristics between the three refractive error groups, the Kruskal-Wallis rank sum test was used for age, Welch's ANOVA for MSFsc, Pearson's Chi-squared test for all other variables.
MSFsc, mid-point of sleep on free days adjusted for sleep debt; Q1, first quartile; Q3, third quartile; SD, standard deviation.

**Table 2.**
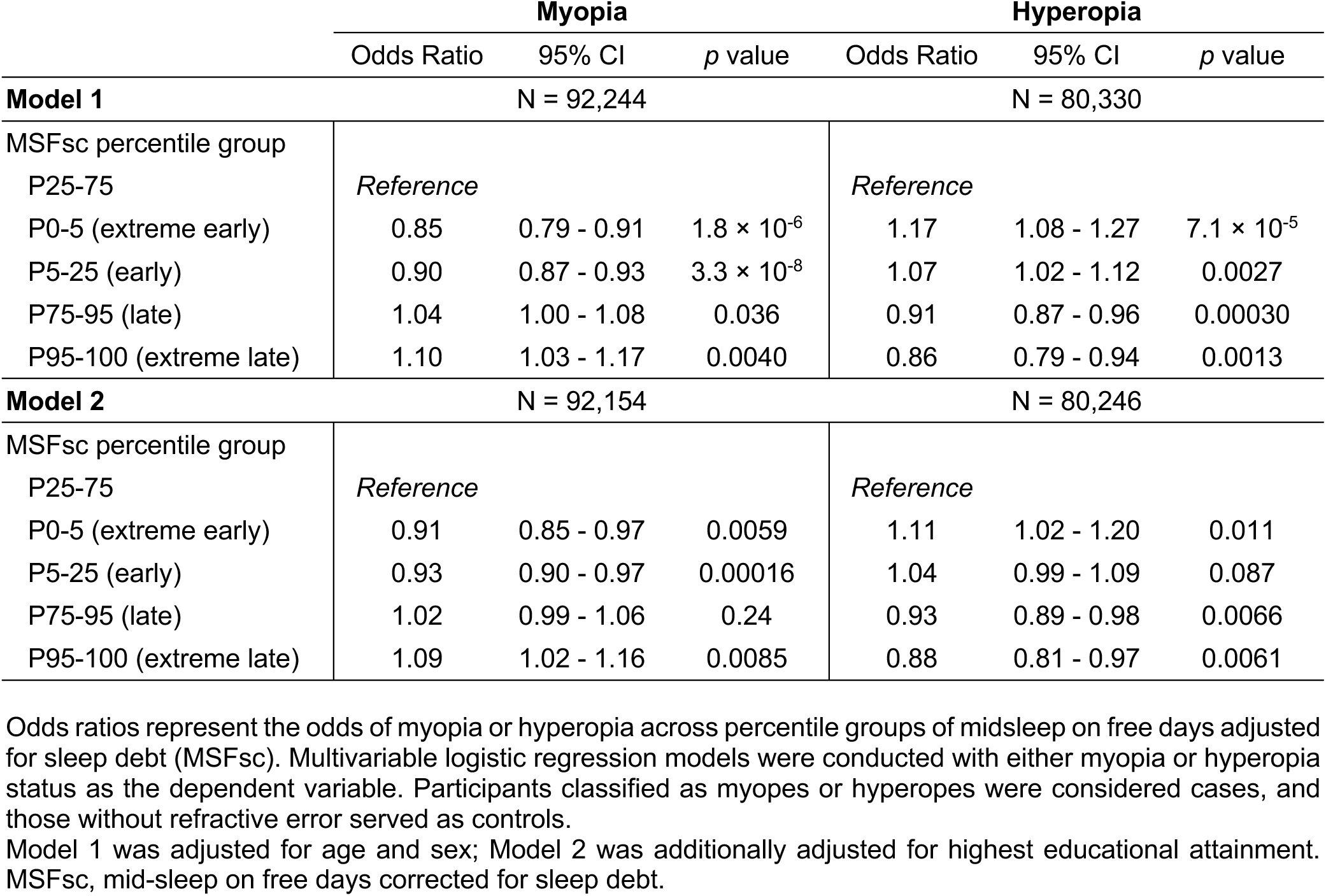
Association of myopia and hyperopia with chronotype in the Estonian Biobank.

To confirm these associations, we conducted replication analyses in the UK Biobank. In this cohort (N = 156,391), 60.7% had no refractive error, 25.9% had myopia, and 13.3% hyperopia (**Table 3**). Chronotype was defined as a three-level variable based on self-reported chronotype (morningness-eveningness: early, intermediate, late). In the age- and sex-adjusted model (**Table 4**, **Model 1**), the UK Biobank reproduced the pattern observed in the Estonian cohort. In the model additionally adjusted for educational attainment (**Table 4**, **Model 2**), again, these associations remained, although modestly attenuated. Compared to morning chronotype, evening chronotype was associated with increased odds of myopia (OR = 1.07, 95% CI: 1.02-1.12, *p* = 0.0048), and reduced odds of hyperopia (OR = 0.92, 95% CI: 0.86-0.98, *p* = 0.0080).

**Table 3.**
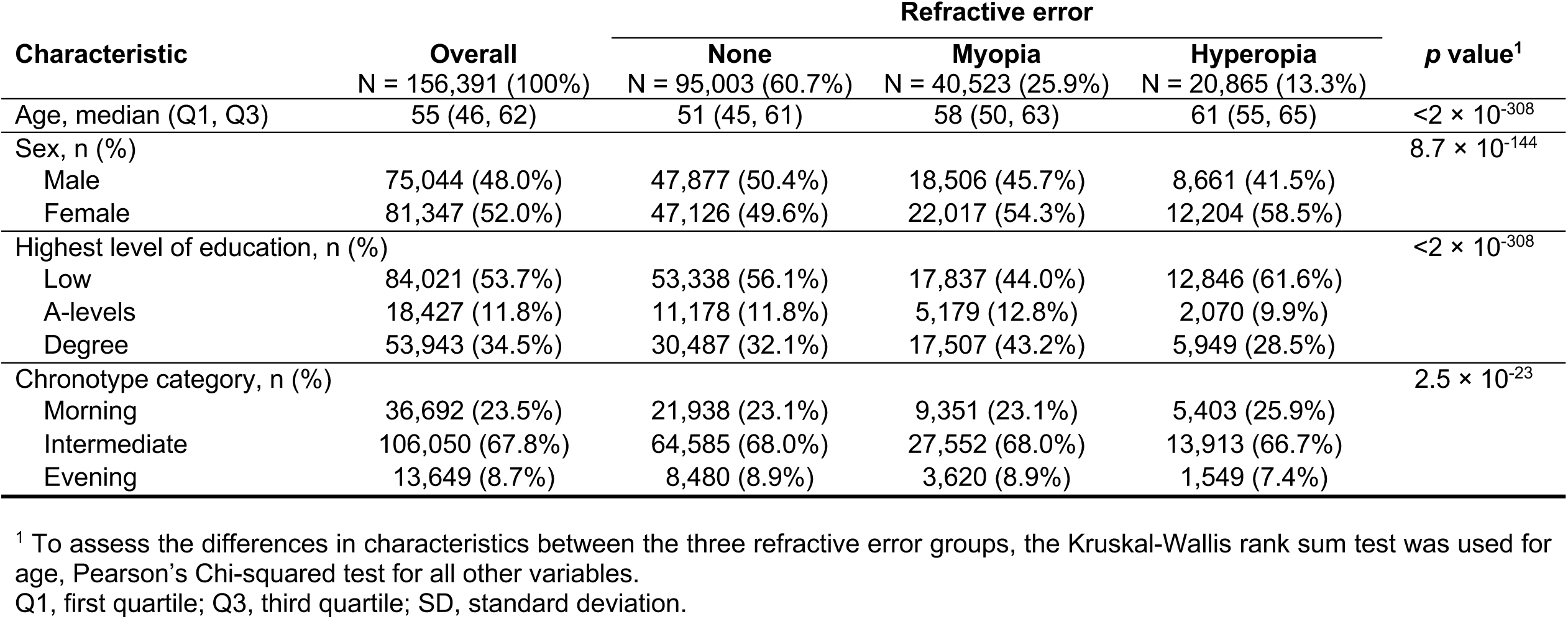
Characteristics of the study population included in the logistic regression analysis from the UK Biobank.

**Table 4.** Association of myopia and hyperopia with chronotype in the UK Biobank.

|  | Myopia |  |  | Hyperopia |  |  |
| --- | --- | --- | --- | --- | --- | --- |
|  | Odds Ratio | 95% CI | *p* value | Odds Ratio | 95% CI | *p* value |
| **Model 1** | N = 137,104 |  |  | N = 117,400 |  |  |
| Chronotype category |  |  |  |  |  |  |
| Early | *Reference* |  |  | *Reference* |  |  |
| Intermediate | 1.08 | 1.04 - 1.11 | $5.5 \times 10^{-7}$ | 0.98 | 0.94 - 1.01 | 0.19 |
| Late | 1.15 | 1.09 - 1.17 | $5.8 \times 10^{-9}$ | 0.91 | 0.86 - 0.98 | 0.0067 |
| **Model 2** | N = 135,526 |  |  | N = 115,868 |  |  |
| Chronotype category |  |  |  |  |  |  |
| Early | *Reference* |  |  | *Reference* |  |  |
| Intermediate | 1.05 | 1.02 - 1.08 | 0.0013 | 0.98 | 0.94 - 1.01 | 0.19 |
| Late | 1.07 | 1.02 - 1.12 | 0.0048 | 0.92 | 0.86 - 0.98 | 0.0080 |

Odds ratios represent the odds of myopia or hyperopia across chronotype categories. Multivariable logistic regression models were conducted with either myopia or hyperopia status as the dependent variable. Participants classified as myopes or hyperopes were considered cases, and those without refractive error served as controls.
Model 1 was adjusted for age and sex; Model 2 was additionally adjusted for highest educational attainment.
CI, confidence interval.

Taken together, these findings demonstrate a robust inverse association between chronotype and refractive error across two large population-based cohorts, with late chronotype linked to a higher likelihood of myopia. Since chronotype reflects the underlying circadian timing of sleep and activity relative to the environment, these correlations raise the possibility that, causally, misalignment between environmental timing cues and the intrinsic clock might influence refractive development.

### Altering the environmental light-dark cycle modulates refractive development in mice

To directly test whether disruption of circadian timing influences refractive development, we altered the environmental light-dark cycle in mice to maintain persistent misalignment between the internal circadian clock and environmental rhythms. C57BL/6J mice (n=13-14 per group) were maintained on a 12 h light / 12 h dark cycle (T24) until postnatal day (P) 28, when baseline ocular measurements were obtained. Animals were then assigned to continued T24 housing or to shortened (T22; 11 h light / 11 h dark) or lengthened (T26; 13 h light / 13 h dark) light-dark cycles (**Fig. 2a** top). Refractive error and ocular biometry were assessed longitudinally. These non-24 h T cycles are a well-established experimental paradigm for challenging the circadian system,^27–29^ requiring repeated daily phase delays (T22) or advances (T26) to maintain entrainment, producing a chronic mismatch between internal circadian timing and the light-dark cycle,^27–29^ analogous to the circadian strain experienced by extreme early and late chronotypes.

**Figure 2.**
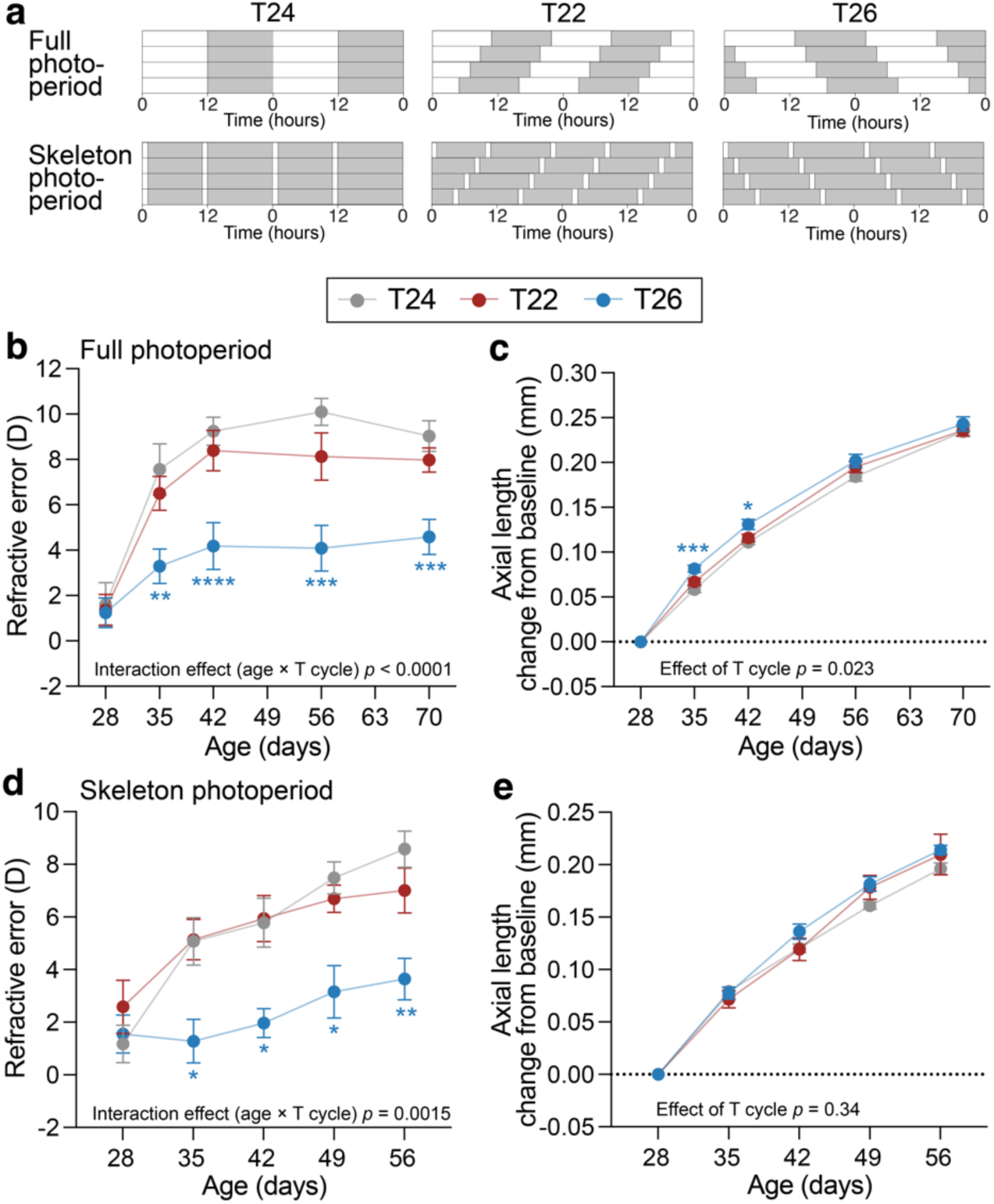
Effects of altered T cycles on refractive development and axial elongation under full and skeleton photoperiods. (a) Experimental light conditions. C57BL/6J mice were maintained under a standard 12 h light-12 h dark cycle (T24) until postnatal day (P)28. At P28, ocular parameters were measured and mice were assigned to one of three T cycle conditions: T24 (12 h light/12 h dark), T22 (11 h light/11 h dark), or T26 (13 h light/13 h dark). Two lighting paradigms were used: full photoperiods (top) and skeleton photoperiods (bottom). In skeleton photoperiods, 1-hour light pulses marked subjective dawn and dusk while maintaining the same T cycle length. Light and dark phases are indicated by white and grey shading, respectively. (b-e) Longitudinal measurements of refractive error (b, d) and axial length change from baseline (c, e) in mice housed in T24, T22, and T26 full photoperiods (b, c) or skeleton photoperiods (d, e). Data are shown as mean ± SEM (full photoperiod: n = 13-14; skeleton photoperiod: n = 6). Statistical analyses were performed using mixed-effects models with Dunnett’s correction for multiple comparisons, testing T22 and T26 against T24 controls. * *p* < 0.05, ** *p* < 0.01, *** *p* < 0.001, **** *p* < 0.0001.

Under standard T24 conditions, mice demonstrated the expected pattern of refractive development, progressing from mild hyperopia at P28 to stable hyperopia by early adulthood (P56 onwards).^30^ Refractive development in T22 did not differ significantly from T24. In contrast, T26 housing altered this trajectory, producing a clear myopic shift relative to controls (**Fig. 2b**; age × T cycle interaction *p* < 0.0001). Although T26-housed mice exhibited a significantly faster rate of axial elongation during the first two weeks, this difference was not sustained at later ages (**Fig. 2c**). Corneal curvature and other ocular biometric parameters, including anterior chamber depth, lens thickness, and vitreous chamber depth, showed no consistent or sustained differences between groups that could account for the more negative refractive error observed in T26-housed mice (**Supplementary Fig 1a-d**).

To isolate the effects of circadian cycle length from light exposure, we used T24, T22, and T26 skeleton photoperiods,^27^ where 1-h light pulses marked subjective dawn and dusk (**Fig. 2a** bottom). Refractive outcomes closely mirrored those observed under full photoperiods: T26 again induced a relative myopic shift, whereas animals housed in T22 showed normal refractive development (**Fig. 2d**; age × T cycle interaction *p* = 0.0015). Axial length (**Fig. 2e**), corneal curvature, and other ocular biometry measures did not differ significantly between groups (**Supplementary Fig. 1e-h**). Visual function and retinal integrity were preserved across all T cycle conditions, indicating that the refractive changes under T26 were not secondary to retinal or visual dysfunction (**Supplementary Fig. 2**).

### Mechanistic insights into the effects of T22 and T26

To characterise the molecular pathways affected by T22 and T26, we performed bulk retinal RNA sequencing after 7 weeks of housing in T22, T24, or T26, with tissue collected at 4 hours after light onset (zeitgeber time, ZT4). In T22, we identified 267 differentially expressed genes (DEGs) relative to T24 (**Fig. 4a**; **Supplementary Table 2**). Among the most significantly upregulated transcripts were several heat shock proteins (*Hspa5*, *Hspa8*), as well as genes involved in photoreceptor differentiation and maintenance, including *Dio2* and the rod-associated transcription factor *Nr2e3*. Gene set enrichment analysis (GSEA) revealed prominent enrichment of synaptic plasticity and neurotransmission pathways, pro-survival and cell adhesion pathways (**Fig. 4d, f**; **Supplementary Table 3**). To quantify cell-type specificity within the adult mouse retina, we analysed the Mouse Retinal Cell Atlas (MRCA), a high-resolution single-cell RNA-seq atlas comprising all major neuronal and non-neuronal retinal classes.^31^ Classifier genes were defined using ROC-based analysis of single-cell transcriptomes, enabling cell-class assignment of bulk DEGs (**Supplementary Fig. 3**, see **Supplementary Methods**). In T22, DEGs mapped across several retinal classes, including glial and vascular populations, but showed no significant enrichment in any single cell type (χ^2^-test, *p* = 0.33; **Fig. 3h** top). Module-score projection to MRCA revealed modest enrichment of up-regulated genes in astrocytes and microglia, while down-regulated genes showed relative enrichment in photoreceptor populations (**Fig. 3i**).

**Figure 3.**
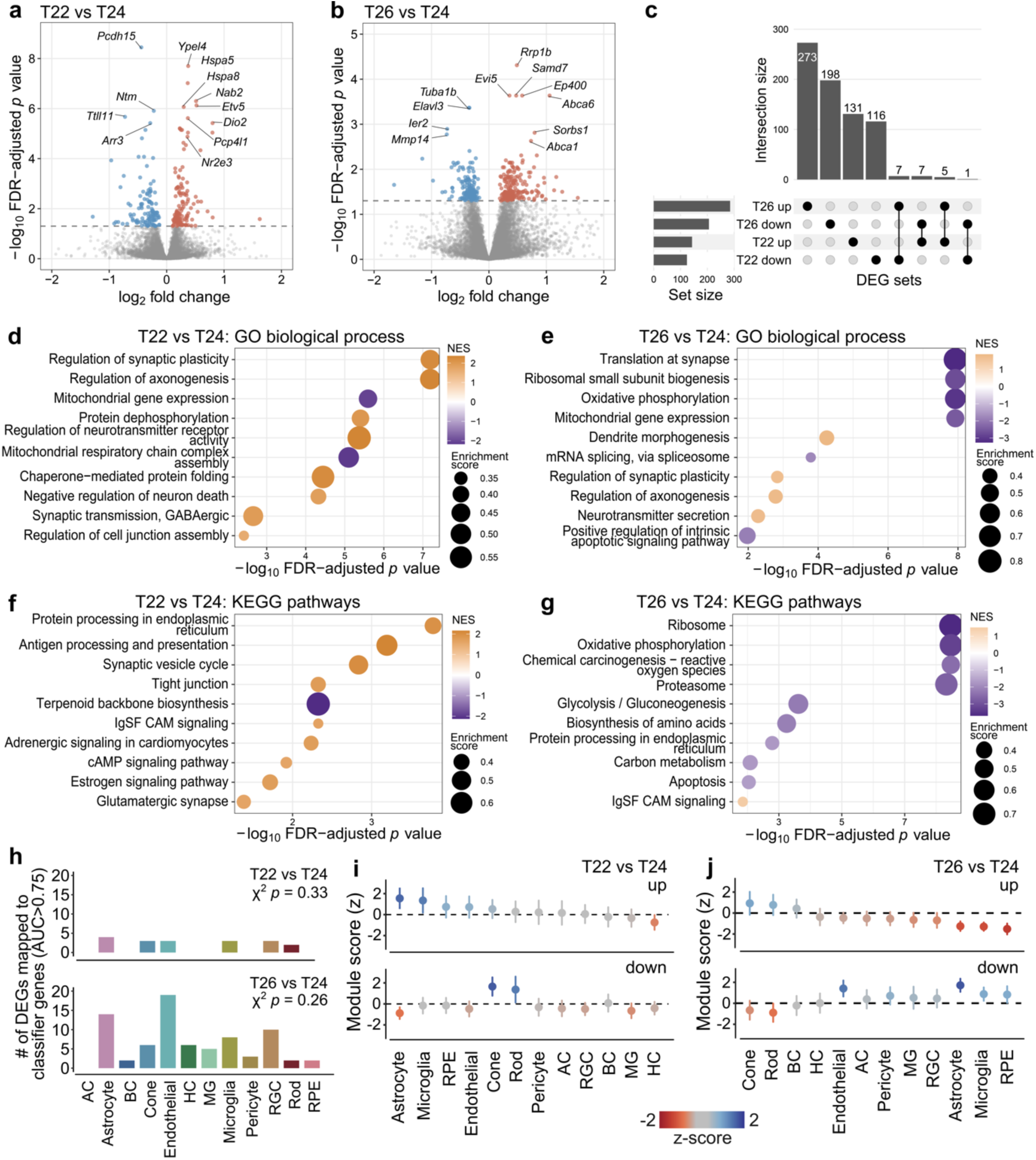
Transcriptional and cell-class-specific responses to altered T cycles in mouse retina. (a-b) Volcano plots of differentially expressed genes (DEGs) for T22 vs T24 (a) and T26 vs T24 (b) after 7 weeks of housing. Selected genes with the lowest adjusted *p* values are labelled. (c) Overlap between upregulated and downregulated DEGs in T22 and T26 relative to T24. (d-g) Gene set enrichment analysis (GSEA) for T22 vs T24 (d) and T26 vs T24 (e), showing selected significantly enriched Gene Ontology (GO) biological process terms (d-e) and KEGG pathways (f-g). Dot colour represents the normalised enrichment score (NES), reflecting direction and magnitude of enrichment, and dot size corresponds to the absolute enrichment score. (h) Mapping of bulk DEGs to major retinal cell classes using classifier genes derived from the Mouse Retinal Cell Atlas (MRCA). Bars indicate the number of DEGs mapping to high-specificity classifier genes (AUC > 0.75) for each retinal class in T22 vs T24 (top) and T26 vs T24 (bottom). χ^2^ test *p* values assess deviation from expected classifier gene frequencies. (i-j) Module-score projection of T22 vs T24 (i) and T26 vs T24 (j) DEGs onto MRCA cell classes. Mean z-scored module values are shown separately for upregulated (top) and downregulated (bottom) gene sets across retinal classes. AC, amacrine cells; BC, bipolar cells; HC, horizontal cells; MG, Müller glia; RGC, retinal ganglion cells; RPE, retinal pigment epithelium.

**Figure 4.**
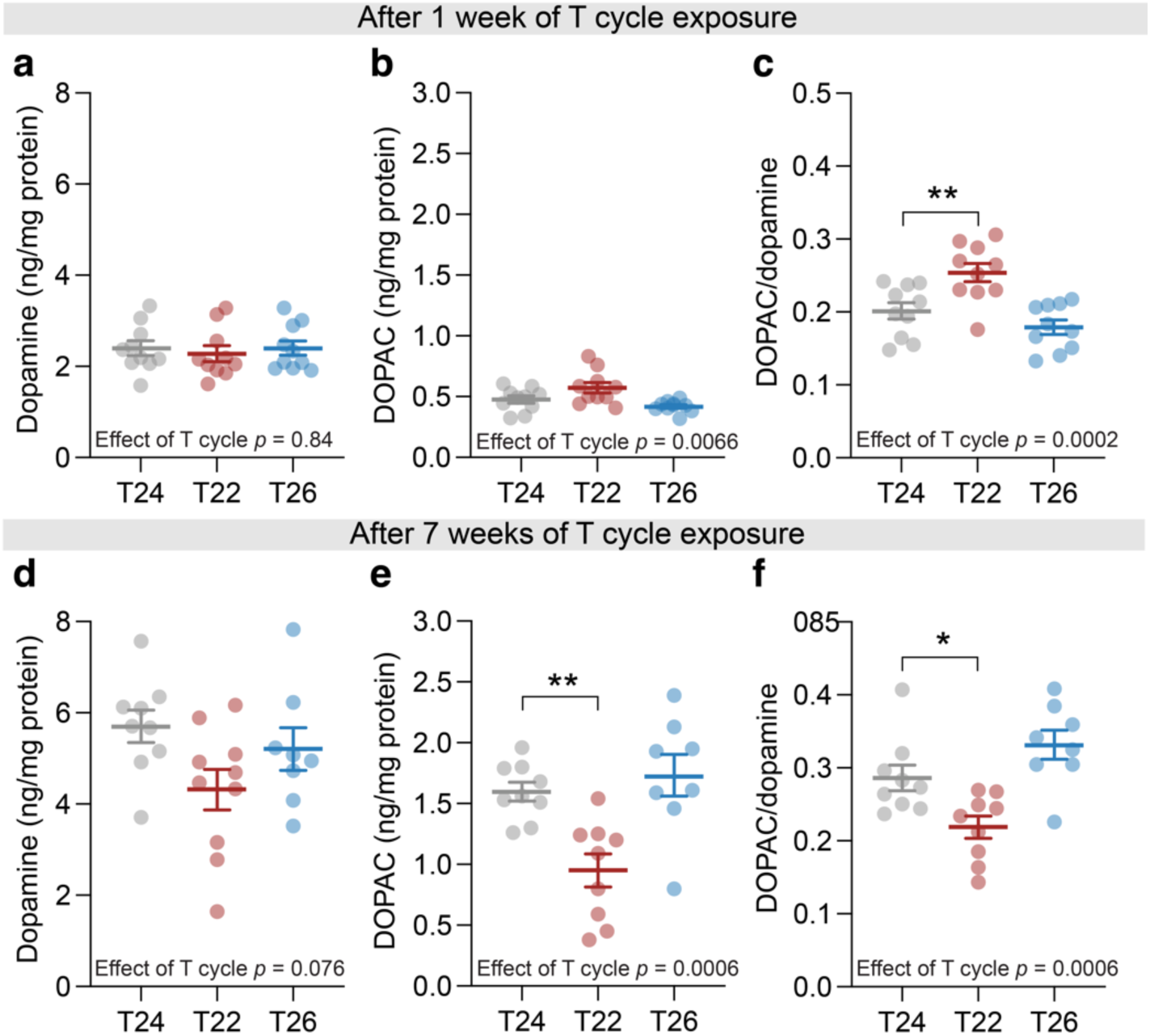
Retinal dopamine and DOPAC levels under altered T cycles. Retinal concentrations of dopamine (a, d), its metabolite DOPAC (b, e), and the DOPAC/dopamine ratio (c, f) after 1 week (a-c) or 7 weeks (d-f) of housing under a standard 12:12-h light-dark cycle (T24), a shortened 11:11-h cycle (T22), or a lengthened 13:13-h cycle (T26). Samples were collected at Zeitgeber time (ZT) 4. Bars show mean ± SEM, with individual data points overlaid, n = 8-10. Statistical analysis was performed using one-way ANOVA with Dunnett’s post hoc test. * *p* < 0.05, ** *p* < 0.01.

In T26, we identified 491 differentially expressed genes relative to T24 (273 upregulated and 198 downregulated; **Fig. 3b**; **Supplementary Table 4**), and there was limited overlap with genes differentially expressed in T22 (**Fig. 3c**). Several of the most significantly altered transcripts were linked to chromatin regulation and transcription (e.g., *Ep400*, *Rrp1b*), cytoskeletal and neuronal structure (*Elavl3*, *Tuba1b*), and lipid transport (*Abca1*, *Abca6*) (**Fig. 3b**). GSEA highlighted strong negative enrichment of mitochondrial oxidative phosphorylation, ribosomal and translational pathways, alongside alterations in carbon metabolism and proteasome function (**Fig. 3e, g**; **Supplementary Table 5**). The DEGs mapped to several retinal cell classes, most prominently astrocytes and endothelial cells, though without significant deviation from expected class distributions (χ^2^-test *p* = 0.26, **Fig. 3h** bottom). Upregulated genes showed relative enrichment in photoreceptor populations, whereas downregulated genes were preferentially expressed in endothelial and glial classes (**Fig. 3j**). Next, to assess whether classical dopamine-mediated pathways, often implicated in myopia,^15,32,33^ could account for the refractive effects observed under altered T cycles, we measured retinal dopamine and its primary metabolite DOPAC at ZT4. After one week of T cycle exposure, dopamine and DOPAC levels were similar between groups, yet dopamine turnover, quantified as the DOPAC/dopamine ratio, was increased in T22 (**Fig. 4a-c**). After 7 weeks of exposure to the different T cycles, DOPAC levels, together with the DOPAC/dopamine ratio, showed a decrease in T22 (**Fig. 4d-f**). Together, these data suggest that dopamine signalling is unlikely to account for the refractive error phenotype observed in T26. The altered dopamine turnover observed in T22 may influence other aspects of retinal physiology but was not associated with measurable changes in refractive development in this study.

### T26 affects refractive development in young adult mice

Finally, we studied whether the effects of T26 on refractive development were confined to early postnatal development. Mice maintained under T24 until P56 and then switched to T26 developed relative myopia compared with animals kept under T24, with an effect size comparable to that observed in juvenile animals (**Fig. 5a**; age × T cycle interaction *p* < 0.0001, n=11-12 per group). Conversely, switching mice from T26 to T24 at P56 reversed the myopic phenotype, indicating that the effects of T26 remain plastic beyond early development. Axial length, other ocular axial parameters and corneal curvature were unaffected by T cycle manipulation (**Fig. 5b**; **Supplementary Fig. 4**).

**Figure 5.**
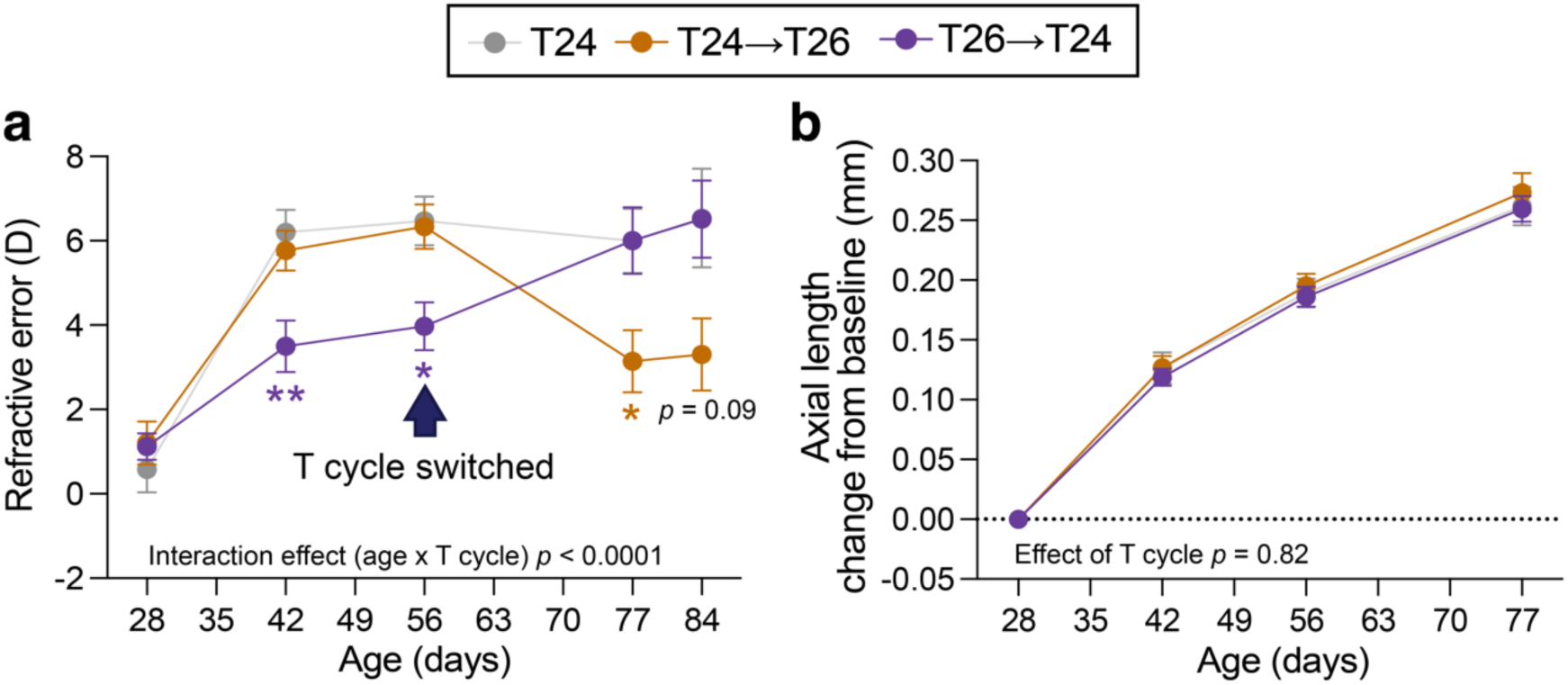
Effects of T26 exposure at adult ages on refractive development. Refractive error trajectories (a) and axial length change from baseline (b) of mice housed under standard 24-h light-dark cycles, mice switched from T24 to T26 at postnatal day (P)56 (T24→T26), and mice switched from T26 to T24 at P56 (T26→T24). Data are mean ± SEM, n = 11-12. Statistical analysis was performed using a mixed-effects model with Dunnett’s correction for multiple comparisons, comparing the intervention groups to T24 controls. * *p* < 0.05, ** *p* < 0.01.

Overall, our findings demonstrate a conserved link between circadian misalignment and refractive development across species. Extending the association between late chronotype and myopia observed in humans, we show that challenging the circadian clock with a lengthened environmental light-dark cycle (T26), but not a shortened cycle (T22), induces a myopic shift in mice. These results support a model in which altered timing between the intrinsic clock and environmental light-dark cycle modulates refractive development, providing a mechanistic link between chronotype and myopia risk.

## Discussion

Refractive errors constitute a growing global health challenge, with myopia alone projected to affect nearly half of the world’s population by 2050,^1,2^ predisposing affected individuals to irreversible sight-threatening ocular diseases and irreversible blindness.^3–6^ Understanding the mechanisms of myopia is, therefore, critical for developing effective prevention and treatment strategies. In two large population-based cohorts from the Estonian and UK Biobanks, with sample sizes of 109,461 and 156,391, respectively, we identified a robust association between late chronotype and myopia. The broad age range also enabled analysis of hyperopia, revealing an opposing pattern whereby early chronotype was associated with increased hyperopia risk. These findings substantially extend prior reports from much smaller cohorts^20,22,21^ and strengthen the evidence for a direction-specific relationship between chronotype and refractive errors.

Our findings in the animal study demonstrate that the misalignment between the intrinsic circadian clock and the external light-dark cycle can directly induce myopia, but the mechanisms linking chronotype and refractive development in humans are likely complex. The rise in myopia prevalence since the mid-20th century parallels major societal changes, including increased educational demands, urbanisation, and widespread use of artificial lighting.^26^ Each of these factors is independently associated with myopia^12,26,34–36^ and late chronotype,^37–43^ suggesting that the observed association between myopia and late chronotype may reflect shared environmental exposures rather than a direct causal relationship.

Light exposure emerges as a particularly compelling mediator. In particular, artificial light at night and evening light exposure, both associated with late chronotype,^41–43^ have also been linked to myopia risk.^35,36^ In addition, reduced exposure to bright outdoor light is a well-established risk factor for myopia^26,44^ and is associated with late chronotype.^17^ Consistent with mediation rather than direct causality, genetic studies have not identified strong genetic correlations or evidence for causal effects between chronotype and myopia.^45,46^ Together, these findings raise the possibility that chronotype may reflect a specific pattern of light exposure and circadian alignment that influences refractive development.

To directly test whether misalignment between the internal circadian clock and the external light environment can drive changes in refractive development, we used a mouse model with non-24-hour light-dark cycles. We found that exposure to a lengthened cycle (T26), but not a shortened cycle (T22) induced myopic eye growth. These findings indicate that the timing of light exposure relative to internal circadian phase, rather than the absolute length of light exposure, is critical. Although T cycles and human chronotype are not equivalent, both late chronotype in humans and T26 entrainment in mice are characterised by short phase angles of entrainment, longer internal circadian periods and light exposure occurring at biologically inappropriate circadian times: during the evening or biological night in humans, and at the onset of the active phase in nocturnal mice.^27,47,16^

The effect of non-24-hour light-dark cycles on ocular phenotypes has, to our knowledge, not been studied. At the molecular level, T22 housing was characterised by synaptic remodelling and pro-survival signalling in the retina, whereas T26 resulted in suppression of mitochondrial oxidative phosphorylation and translational pathways, consistent with activation of hypoxia-related pathways that could contribute to altered refractive development and have previously been implicated in experimental myopia.^48^ Across both conditions, transcriptional changes were broadly distributed across retinal classes, with relatively stronger module-score enrichment in photoreceptors and astrocytes, indicating a multicellular response rather than a single cell-type-restricted driver.

Furthermore, we show that T26-induced myopic shifts are not confined to early postnatal development but can also be induced and reversed in young adult mice. This contrasts with conventional myopia paradigms, in which refractive plasticity is expected to decline sharply after early adolescence. In a single study investigating experimental myopia induction in mice at later ages, form-deprivation, another experimental myopia model, induced axial elongation as late as P40, but plasticity was abolished by P67.^49^ Whether the phenomenon of adult plasticity is unique to circadian manipulation or reflects a broader capacity of the adult eye to respond to altered temporal or visual inputs warrants further investigation.

Besides the novel insights, several limitations should be considered. In the human cohorts, chronotype was assessed in adulthood, whereas refractive development largely occurs during childhood and adolescence, limiting temporal inference. Both biobanks were predominantly of European ancestry,^50,51^ and refractive status was derived from health records or self-report rather than from direct measurement of refractive error. In the mouse experiments, we could not identify a single biometric parameter fully accounting for the refractive changes; however, the expected axial length differences in mice are extremely small and often fall below reliable detection limits.^52,53^ Future studies incorporating longitudinal refractive error measurements, objective light exposure and sleep assessments in humans, more diverse populations, and higher-resolution ocular and molecular analyses in animal models, will be important to refine these findings.

In conclusion, our study identifies circadian misalignment as a conserved modulator of refractive development and suggests that the timing of light exposure relative to the internal circadian state is a potentially modifiable risk factor. These findings have important public health implications, raising the possibility that strategies promoting appropriate light exposure and circadian-aligned sleep-wake behaviours, alongside existing recommendations for outdoor time, may represent accessible, low-cost approaches to mitigating myopia risk in modern environments.

## Methods

### The Estonian Biobank cohort and phenotyping

The Estonian Biobank is a population-based cohort of adult volunteers (N=211,979) with linked health-registry data. The population is described in full elsewhere.^50,54^ All participants provided broad informed consent. Analyses were conducted in accordance with the Human Genes Research Act,^55^ with approval from the Estonian Committee on Bioethics and Human Research (approval 1.1-12/624), and the Estonian Biobank data release application 3-10/GI/34223. Chronotype was assessed using the Estonian translation^56^ of the Munich Chronotype Questionnaire (MCTQ) and quantified as midsleep on free days corrected for sleep debt (MSFsc). To account for the strong age dependence of MSFsc, participants were grouped into age-specific chronotype percentile categories. Myopia and hyperopia were defined using a combination of ICD-10 diagnoses and self-reported refractive status. Regression models were adjusted for age, sex, and educational attainment. Full details of cohort characteristics, phenotyping, quality control, and covariate definitions are provided in **Supplementary Methods**.

### The UK Biobank cohort and phenotyping

The UK Biobank is a prospective general population cohort of over 502,000 UK residents. The study population is described in full elsewhere.^57,58^ The UK Biobank has received ethical approval from the North West Multi-Centre Research Ethics Committee (11/NW/03820), and participants have provided written, informed consent. Chronotype was defined using a self-reported morningness-eveningness question and categorised as early, intermediate, or late. Myopia and hyperopia were defined based on self-reported use of glasses or contact lenses for refractive correction. Analyses were adjusted for age, sex, and educational attainment (see **Supplementary Methods**).

### Statistical analysis of epidemiological data

Associations between chronotype and refractive error were assessed using logistic regression models, with myopia or hyperopia as the outcome and participants without refractive error as controls. Models were adjusted for age and sex, with additional adjustment for educational attainment (see **Supplementary Methods**).

### Animal experiments

All animal procedures conformed to the ARVO Statement for the Use of Animals in Ophthalmic and Vision Research and were approved by the Emory University Institutional Animal Care and Use Committee (IACUC; protocol PROTO202300061). C57BL/6J mice of both sexes were used. Mice were housed under controlled light-dark conditions with food and water provided *ad libitum*.

To challenge circadian timing, mice were transferred to a shortened (T22; 11 h light / 11 h dark) or lengthened (T26; 13 h light / 13 h dark) light-dark cycle from P28. Additional experiments used skeleton photoperiods to dissociate circadian cycle length from total light exposure, and young adult mice (P56) were switched between T24 and T26 schedules to assess reversibility of the phenotype. Refractive error and ocular biometry were assessed longitudinally.^59^ Retinal function was assessed using electroretinography, visual function with the optomotor reflex, and retinal structural integrity by histology. Dopamine and its primary metabolite DOPAC levels were quantified using high-performance liquid chromatography, and dopamine turnover was estimated using the DOPAC/dopamine ratio.^60^

Statistical analyses of animal experiments described above were performed using GraphPad Prism (v10). Data are presented as mean ± SEM. Group comparisons were conducted using one- or two-way ANOVA with corrections for multiple comparisons, and mixed-effects models for longitudinal data with missing values. The specific tests used are indicated in the figure legends, and *p* < 0.05 was considered statistically significant.

### RNA-sequencing procedure and analysis

Retinal transcriptomic profiling was performed using bulk RNA sequencing. Total RNA was extracted from retinas collected at ZT4, quality-controlled using Agilent 2100 Bioanalyzer (Agilent Technologies), and samples with an RNA integrity number (RIN) >9.0 were used. Libraries were prepared using poly(A)-selection, followed by paired-end sequencing on the Illumina platform. Low-quality reads and adapters were removed using fastp,^61^ and read quality was assessed with FastQC^62^ (see **Supplementary Table 6** for RNA and sequencing quality control measures). Transcript abundance was quantified against the mouse reference genome using Salmon,^63^ and summarised at the gene level. Differential expression analysis was performed using DESeq2,^64^ with adjustment for sex and circadian phase.^65^ To identify affected biological pathways, gene set enrichment analysis was performed using clusterProfiler^66^ against Kyoto Encyclopedia of Genes and Genomes (KEGG) pathways^67^ and Gene Ontology (GO) biological process categories.^68^ To estimate cell-type contributions in bulk retinal RNA-seq data, deconvolution analysis was performed by integrating bulk differentially expressed genes with the Mouse Retinal Cell Atlas^31^ using Seurat (v4.0).^69^ Cell-type-specific classifier genes were identified using receiver operating characteristic (ROC) analysis, and enrichment of bulk transcriptional signatures across retinal classes was quantified by module scoring. Full details are provided in the **Supplementary Methods**. The RNA-sequencing data (raw reads and processed gene-level matrices) will be made publicly available via NCBI’s Gene Expression Omnibus (GEO) upon publication.

## Acknowledgments

We want to acknowledge the participants of the Estonian Biobank for their contributions. This research was supported by the European Union through the Horizon 2020 research and innovation program under grant no. 894987, by the European Regional Development Fund project no. MOBEC008, the Estonian Research Council grant PRG1291, and the European Union’s Horizon Europe MSCA fellowship grant No. 101153901. Views and opinions expressed are however those of the authors only and do not necessarily reflect those of the European Union or European Research Executive Agency (REA). Neither the European Union nor the granting authority can be held responsible for them. The research was conducted using the Estonian Centre of Genomics/Roadmap II, funded by the Estonian Research Council (project number TT17). This work was also supported by NIH R01 EY016435, NIH R01 EY033361, the Department of Veterans Affairs Research Career Scientist Award RX003134, the Research to Prevent Blindness Challenge Award, NIH P30EY006360, and NIH R01 HG012810. The study was supported in part by the Emory Integrated Genomics Core (EIGC), which is subsidised by the Emory University School of Medicine and is one of the Emory Integrated Core Facilities. The study was supported in part by the Emory HPLC Bioanalytical Core (EHBC), which was supported by Emory University School of Medicine and the Georgia Clinical & Translational Science Alliance of the National Institutes of Health under Award Number UL1TR002378. The content is solely the responsibility of the authors and does not necessarily reflect the official views of the National Institutes of Health. This work was partly written at writing retreats and writing days organised by the Institute of Genomics, University of Tartu. The Estonian Biobank Research Team was responsible for data collection in the Estonian Biobank and consisted of Andres Metspalu, Mait Metspalu and Lili Milani.

## Author contributions

T.P. and E.A. conceived and designed the study. T.P. and M.T.P. designed the animal experiments. T.P., N.T. and M.T.-L. performed the MCTQ calculations and data cleaning. T.P. and N.T. performed the sample characterisation and statistical analyses in the Estonian Biobank cohort. A.C.B. and J.M.C. performed all analyses in the UK Biobank cohort. T.P. and S.B. performed the animal experiments. S.P.D. performed the single-cell-informed cell-type deconvolution analyses. R.S., T.E, E.A., M.T.P., and P.P. supervised the study. T.P. wrote the first draft of the manuscript. All authors critically reviewed the manuscript. M.T.P and P.P. contributed equally to this work.

## Data availability

Pseudonymised genotype and phenotype data are available from the Estonian Biobank (https://genomics.ut.ee/en/content/estonian-biobank) upon request. The RNA-sequencing data (raw reads and processed gene-level matrices) will be made publicly available via NCBI’s Gene Expression Omnibus (GEO) upon publication.

## Code availability

Analyses used are described in the Methods and Supplementary Methods sections of the manuscript

