## Supplementary Information for "Circadian rhythms regulate refractive development across species"

This PDF file includes:

Supplementary Methods
Supplementary Figures 1-4
Supplementary Table 1

Other Supplementary Information for this manuscript includes the following:

Supplementary Tables 2-6, provided as an Excel file

#### Supplementary Methods

##### The Estonian Biobank

###### Data and ethics

The Estonian Biobank is a population-based, volunteer cohort comprising approximately 211,979 adult participants.[1,2] All participants have provided broad informed consent. Health-related information, including medical diagnoses, is obtained through regular linkage with national registries, primarily the Estonian National Health Insurance Fund, and diagnoses are coded according to the International Classification of Diseases, 10th revision (ICD-10). The majority of electronic health records have been collected since 2004.[1] The activities of the Estonian Biobank are governed by the Human Genes Research Act.[3] Individual-level data analyses were conducted under approval 1.1-12/624 from the Estonian Committee on Bioethics and Human Research (Estonian Ministry of Social Affairs), and in accordance with the approved Estonian Biobank data release application 3-10/GI/34223.

###### Chronotype definition

Sleep timing and chronotype were assessed using the Estonian translation[4] of the Munich Chronotype Questionnaire (MCTQ),[5] which is included in the standard questionnaire administered to Estonian Biobank participants. The MCTQ collects self-reported sleep timing information separately for workdays and non-workdays over the preceding 4 weeks. Participants reporting shift work and those with extreme or implausible values for sleep timing or sleep duration were excluded according to predefined quality-control criteria (**Supplementary Table 1**). Chronotype was quantified as midsleep time on free days corrected for sleep debt (MSFsc). Because MSFsc shows a strong non-linear relationship with age, participants were stratified into age-specific MSFsc percentile groups to allow comparison across relative chronotype categories. Within each 1-year age bin, the 5th, 25th, 75th, and 95th percentiles of the MSFsc distribution were calculated. Based on these cut-offs, individuals were classified as extreme early (P0-5), early (P5-25), intermediate (P25-75), late (P75-95), or extreme late (P95-100) chronotypes. The intermediate group (P25-75) was used as the reference category in regression analyses.

Myopia and hyperopia definitions

Participants were classified as myopia cases if they had an ICD-10 diagnosis of myopia (H52.1) recorded in health registers or if they self-reported a diagnosis of myopia. Hyperopia cases were defined analogously using ICD-10 code H52.0 or self-reported hyperopia. Individuals with records indicating both myopia and hyperopia were excluded from the analysis. All remaining participants were treated as controls. Sample counts and exclusions are provided in **Supplementary Table 1**.

Covariates

The highest level of education was self-reported on a nine-level scale and grouped into three categories: basic education (no primary education, primary education, or basic education), secondary education (secondary or vocational secondary education), and higher education (applied higher education, university degree, or advanced academic degree). Age and education were defined at the time of MCTQ completion. Participants aged 80 years or older were excluded from analyses, as age-related ocular and neurological conditions can influence sleep patterns and circadian regulation.[6,7]

#### The UK Biobank

Data and ethics

The UK Biobank is a prospective general population cohort, which contains more than 502,000 UK residents recruited via National Health Service patient registers from 2006 to 2010. The study population is described in full elsewhere.[8,9] The UK Biobank has received ethical approval from the North West Multi-Centre Research Ethics Committee (ref 11/NW/03820). Participants who accepted the invitation to join the UK Biobank cohort provided written, informed consent.

Chronotype definition

Chronotype assignment was based on a single self-reported measure (“Morning/evening person (chronotype)”, UK Biobank data field 1180). Participants answered the question “Do you consider yourself to be?” by selecting one of six response options. Participants reporting “definitely a ‘morning’ person” were classified as early chronotype, and those reporting “definitely an ‘evening’ person” were classified as late chronotype. Participants selecting either intermediate response (“more a ‘morning’ than ‘evening’ person”, “more an ‘evening’ than a ‘morning’ person”) were grouped as intermediate chronotype. Individuals responding “do not know” or “prefer not to answer” were excluded. Early chronotype was used as the reference category in all analyses.

Myopia and hyperopia definitions

Refractive error assignment was based on the UK Biobank self-reported data fields 2207 and 6147. For the question “Do you wear glasses or contact lenses to correct your vision?” (data field 2207), individuals who answered “No” were used as controls. A “Yes” answer prompted the follow-up question “Why were you prescribed glasses/contacts? (You can select more than one answer)” (data field 6147), which includes the options “For short-sightedness, i.e. only or mainly for distance viewing such as driving, cinema etc (called ‘myopia’)” and “For long-sightedness, i.e. for distance and near, but particular for near tasks like reading (called ‘hypermetropia’)”. Individuals who responded with “For short-sightedness” while not responding with “For long-sightedness” were included as cases for myopia. Individuals who included the response “For long-sightedness” and not “For short-sightedness” were included as cases of hyperopia. Additionally, individuals who answered the question “Why were you prescribed glasses/contacts? (You can select more than one answer)” with options other than hyperopia or myopia were also used as controls.

Covariates

The highest level of education was self-reported in the UK Biobank data field 6138, which records participants’ highest educational qualification across seven predefined categories. These were further grouped into three: Degree (College or University degree), A-levels (A levels / AS levels), Low (O levels / GCSEs, CSEs, NVQ / HND / HNC, other professional qualifications). Age at replying to the sleep habit questionnaire is reported in the article.

#### Statistical analysis of epidemiological data

To assess the association between chronotype and myopia/hyperopia, we fitted separate logistic regression models with myopia or hyperopia status as the dependent variable, including participants classified as myopes or hyperopes, respectively, as cases, and participants with no refractive error as controls. The models were adjusted for age and sex and additionally for education level. Multicollinearity was tested by calculating the variance inflation factor for independent variables in regression models, and no multicollinearity was detected between the variables of interest.

#### Animal experiments

##### Animal husbandry and housing

All procedures conformed to the ARVO Statement for the Use of Animals in Ophthalmic and Vision Research and were approved by the Emory University Institutional Animal Care and Use Committee (protocol PROTO202300061). C57BL/6J mice (The Jackson Laboratory, strain #000664; Bar Harbor, ME, USA) were used. Mixed cohorts of male and female mice were included. The number of animals in each experiment is noted in the figure legends. Animals were delivered on postnatal day 21 (P21), acclimated for 1 week, and experimental procedures began at P28. Mice were group-housed under overhead white light-emitting diode (LED) illumination (100 lux, or 14.1 log photon flux at cage floor) with water and standard rodent chow available *ad libitum*. Animals were randomly assigned to experimental conditions.

##### Experimental procedure

Mice were housed under a standard 12:12-h light-dark cycle (T24) from birth until postnatal day 28 (P28). At P28, baseline ocular parameters were measured, including refractive error, corneal curvature and ocular biometry. Mice were then randomly assigned to one of three conditions: T24: 12h light / 12h dark (control), T22: 11h light / 11h dark, T26: 13h light / 13h dark. In a separate set of experiments, skeleton photoperiods were used to dissociate the effects of photoperiod duration from circadian cycle length. In these conditions, mice were housed under T22 or T26 skeleton schedules consisting of two 1-h light pulses marking subjective dawn and dusk. In another set of experiments, mice were maintained under a standard 12 h light / 12 h dark cycle until postnatal P56, and then transferred to a T26 light-dark cycle. A complementary group was switched from T26 back to T24 at P56 to assess reversibility of the phenotype.

For all experiments, ocular measurements were repeated weekly or biweekly throughout the experimental period, as indicated for each experiment. At the end of the study, visual function was assessed by optomotor reflex (OMR), retinal function by ERG and eyes were collected for retinal transcriptomics and monoamine analysis. The experimental groups consisted of 6-14 animals per group per experiment, an equal mix of male and female mice; exact sample sizes are specified in the corresponding figure legends.

Refraction, corneal curvature, and ocular biometry measurements

Refractive error was measured using an infrared automated photorefractor.[10] Following pupil dilation with 1% tropicamide and confirmation of full dilation, mice were anaesthetised via intraperitoneal injection of ketamine (100 mg/kg) and xylazine (10 mg/kg). Measurements were obtained nasally or temporally, just outside the optic nerve head, along the optical axis. Anterior corneal radius of curvature was then assessed using an automated keratometer. Biometric measurements were acquired with a 1310 nm spectral-domain optical coherence tomography (SD-OCT) system (Envisu R4300; Bioptigen, Durham, NC, USA). Three whole-eye OCT scans were obtained 10° superior to the optic nerve head, and ocular tissue interfaces were manually marked to derive anterior chamber depth (ACD), lens thickness (LT), vitreous chamber depth (VCD), and axial length (AL). Axial length was defined as the distance from the anterior corneal surface to the anterior border of the retinal pigment epithelium. Following all measurements, anaesthesia was reversed with atipamezole (0.5 mg/kg), and mice were monitored on a warming pad until full recovery.

Electroretinography

Following overnight dark adaptation, mice were anaesthetised with ketamine (100 mg/kg) and xylazine (10 mg/kg). All procedures were conducted under dim red light. Pupils were dilated with 1% tropicamide (Alcon Laboratories), and corneal analgesia was induced with proparacaine (Alcon Laboratories). Body temperature was maintained at 37°C throughout using a heating pad. Recordings were obtained using a commercial system (UTAS 3000; LKC Technologies, Gaithersburg, MD, USA) with Ganzfeld stimulation. Full-field flash stimuli were presented under dark-adapted conditions, with flash intensities and interstimulus intervals progressively adjusted to span the dynamic range of light responses. Gold-loop electrodes were placed on the corneal surface through a layer of 0.5% methylcellulose, and reference electrodes (platinum subdermal needles; Natus Medical, Pleasanton, CA, USA) were inserted subcutaneously in the cheeks, with a common ground electrode in the tail. After recordings, anaesthesia was reversed with atipamezole (0.5 mg/kg), lubricating eye drops were applied, and mice were allowed to recover on a heating pad. The a-wave amplitude was measured from baseline to the first negative trough. The b-wave amplitude was measured from the a-wave trough (or baseline if absent) to the positive peak. Responses were averaged from both eyes.

Visual function

Visual acuity and contrast sensitivity were assessed using the OptoMotry system[62] (CerebralMechanics, Lethbridge, AB, Canada), which measures optokinetic tracking responses to rotating sine wave gratings. Mice were placed unrestrained on a platform at the centre of a virtual cylinder formed by computer monitors. Visual acuity (spatial frequency threshold) and contrast sensitivity were determined using the system's "Auto Advise" mode, which employs a stepwise staircase protocol. In this protocol, stimulus parameters (spatial frequency or contrast) were automatically increased if the mouse responded and decreased if the mouse did not respond, producing a back-and-forth sequence. After seven such reversals, the software automatically calculated the threshold as the mean of the final three reversals. Measurements were performed under photopic conditions, and testing was completed for clockwise and anti-clockwise rotations to stimulate each eye independently. The average value from both eyes was used for analysis.

HPLC analysis of monoamines

Retinal samples were collected, immediately frozen on dry ice, and stored at −80 °C until analysis. For high-performance liquid chromatography (HPLC), 100 µL of 0.1 M perchloric acid was added to each sample, followed by sonication for 10 s at a 30% duty cycle. Homogenates were centrifuged (10 min, 10,000 × g, 4 °C), and supernatants were transferred to 0.22 µm PVDF filter tubes. Pellets were stored at −80 °C for protein analysis, which was performed using the Pierce™ BCA Protein Assay Kit (Thermo Fisher Scientific, Cat. No. 23225). Filters were centrifuged at 5,000 × g for 2 min, and filtrates were submitted to the Emory HPLC Bioanalytical Core for monoamine analysis. Subsequently, monoamines were quantified using an ACQUITY ARC system (Waters) with a 3465 electrochemical detector. Separations were performed at 37 °C on an XBridge BEH C18 column (2.5 µm, 3 × 150 mm; Waters) with a mobile phase of 100 mM citric acid, 100 mM phosphoric acid, 0.1 mM EDTA, 525 mg/L OSA, and 7% acetonitrile (pH 3.3) at a flow rate of 0.6 mL/min. A 20 µL sample was injected, with water as the needle wash. Pump piston wash was 15% isopropanol. Detection used a SenCell flow cell (2 mm GC WE) with a cell potential of 800 mV (salt bridge reference electrode), automatic step time (AST) position 1, and analogue digital filter (ADF) set to 0.5 Hz. Monoamine concentrations were normalised to protein content and expressed as nanograms per milligram of protein.

Histology and marking retinal layers

Eyes were enucleated and fixed in 97% methanol and 3% glacial acetic acid at −80°C for at least four days. Fixed tissues were embedded in paraffin, and 5 µm sections were prepared through the optic nerve head (ONH) in the cross-sectional plane. Sections were deparaffinised and rehydrated using a graded xylene-ethanol series, followed by standard haematoxylin and eosin staining. Sections were imaged by brightfield microscopy (Nikon Ti, DS-Ri2 camera, 20× objective; 0.37 µm/pixel). Retinas were assessed at regularly spaced intervals of 500 µm apart. Outer nuclear layer nuclei were manually counted in a 100 µm-wide section using the ImageJ cell counter plugin.

RNA extraction and sequencing

Total RNA was extracted from retinas using QIAzol reagent and the RNeasy Micro Kit (QIAGEN). Eyes were enucleated, immediately flash-frozen on dry ice, and stored at −80 °C until processing. Retinas were dissected in ice-cold PBS and homogenised in 400 µL QIAzol using a tissue homogeniser (Argos Technologies). Homogenates were incubated at room temperature for 5 min, placed on ice, and then mixed with 100 µL chloroform (Sigma-Aldrich) by vigorous shaking for 15 s. Samples were incubated at room temperature for 3 min, returned to ice, and centrifuged at 12,000 × g for 15 min at 4 °C. The upper aqueous phase (~200 µL) was transferred to a new nuclease-free tube, mixed with an equal volume of 70% ethanol, and loaded onto RNeasy MiniElute spin columns. RNA purification was completed following the RNeasy Micro Kit protocol provided by the manufacturer.

RNA quality was assessed using the Agilent 2100 Bioanalyzer (Agilent Technologies) with an RNA 6000 Pico Kit at the Emory Integrated Genomics Core. Samples with an RNA integrity number (RIN) >9.0 were considered suitable for RNA sequencing. RIN values of these samples ranged from 9.1 to 9.6. Samples were submitted to Admera Health for further library preparation and sequencing. Libraries were prepared using the NEBNext Ultra II Directional kit with Poly-A selection and sequenced at a depth of 40 million total reads per sample, 2 × 150 bp configuration, on the Illumina platform.

RNA-seq data analysis

Low-quality reads and adapters were removed using fastp,[12] using default parameters, and poly-X tail trimming was enabled to account for the poly(A)-enriched library preparation. Read quality was assessed with FastQC.[13] See RNA, sequencing, and read quality post-trimming in **Supplementary Table 6**. Transcript abundance was quantified with Salmon,[14] with decoy-aware selective alignment. A selective-alignment index was constructed from the *Mus musculus* reference genome (GRCm39) and Ensembl transcriptome (release 113). Quantification was performed with --validateMappings and default alignment scoring parameters. Transcript-level abundance estimates were summarised to the gene level prior to analysis.

Count normalisation and differential expression analysis were performed using DESeq2.[15] Lowly expressed genes were filtered prior to differential expression analysis by retaining only genes with at least 10 counts in at least six samples (corresponding to the smallest group size). To account for inter-individual differences in circadian state despite standardised tissue collection at zeitgeber time 4 (ZT4), we inferred a relative “clock state” from core clock gene expression, adapting an approach described by Talamanca *et al.*[16] From a canonical set of mammalian clock genes, we selected genes with sufficient expression variability in our dataset (median absolute deviation ≥ 0.10 in variance-stabilised expression space), yielding six genes (*Ciart, Dbp, Per1, Per2, Nr1d1, Nr1d2*). Principal component analysis (PCA) was performed on the expression of these genes across all samples. Samples were projected into PC1-PC2 space, and a circular clock state variable (θ) was derived from the angular position using the two-argument arctangent function. Sine and cosine transformations of θ were included as covariates in an extended DESeq2 model: design = ~ cos(θ) + sin(θ) + Sex + Treatment. The RNA-sequencing data (raw reads and processed gene-level matrices) will be made publicly available via NCBI's Gene Expression Omnibus (GEO) upon publication.

Gene set enrichment analysis (GSEA)

GSEA was performed using clusterProfiler,[17] with genes ranked by the DESeq2 Wald statistic. Enrichment was assessed against Kyoto Encyclopedia of Genes and Genomes (KEGG) pathway annotations[18] and Gene Ontology (GO) biological process categories.[19] Statistical significance was determined using Benjamini-Hochberg false discovery rate (FDR) correction, with FDR-adjusted $p$ value < 0.05 considered significant. For each gene set, an enrichment score (ES) was calculated using a running-sum statistic that increases when a gene in the ranked list belongs to the gene set and decreases otherwise, reflecting the degree to which genes from the set are overrepresented at the top or bottom of the ranked list.[17] The normalised enrichment score (NES) was obtained by dividing the observed ES by the mean absolute ES from permutation testing, thereby normalising for gene set size and enabling comparison across gene sets. Positive NES values indicate enrichment among genes upregulated in the comparison group, whereas negative NES values indicate enrichment among downregulated genes.

Mouse Retinal Cell Atlas (MRCA) processing

We analysed the Mouse Retinal Cell Atlas (MRCA),[20] an integrated single-cell RNA-sequencing dataset encompassing major retinal neuronal and non-neuronal classes. Initial processing followed the Seurat (v4.0) workflow.[21] Raw count matrices stored in Ensembl gene nomenclature were first matched to an accompanying gene-ID conversion table. Ensembl identifiers with missing or duplicated gene names were removed, and unique gene symbols were enforced using make.unique(). The resulting matrix was subset to genes present in both the raw assay and the annotation table, and row names were replaced with gene symbols.

A Seurat object was created using CreateSeuratObject (min.features = 200, min.cells = 3) while preserving the original metadata. Standard quality-control metrics were computed, including total UMI counts, number of detected genes, and mitochondrial transcript percentage (PercentageFeatureSet, pattern = "^mt-"). Cells were retained if they met the following criteria: >200 detected genes, <7000 detected genes, <35,000 UMIs, and <12% mitochondrial reads. Data were normalised with NormalizeData, and 3000 highly variable genes were identified using the "vst" method (FindVariableFeatures). Counts were scaled while regressing out mitochondrial percentage (ScaleData).

Dimensionality reduction was performed using principal component analysis (RunPCA, 50 PCs), and uniform manifold approximation and projection (UMAP) embeddings were generated using the first 30 PCs (RunUMAP). Major retinal classes, including rods, cones, bipolar cells, Müller glia, amacrine cells, horizontal cells, retinal ganglion cells, endothelial cells, pericytes, astrocytes, microglia, and retinal pigment epithelium, were used to define cell identities (SetIdent). Quality of annotation was confirmed using known markers (e.g., *Six3*) and by evaluating UMAP distributions grouped by ontology labels and major cell classes. The processed atlas was used for all downstream analyses.

ROC analysis and cell-type specificity assessment

To identify genes with high specificity for each retinal cell class, we applied a receiver-operator characteristic (ROC)-based differential expression approach implemented in Seurat's FindMarkers with test.use = "roc". For each major cell class, the classifier compared cells of the target identity (ident.1) against all other cells in the atlas, capped at 1000 cells per identity to standardise representation. ROC tests report the area under the ROC curve (AUC; "myAUC") as a measure of discriminatory power, with AUC values approaching 1 indicating high class specificity. ROC analyses were run for all major retinal classes, and results for each class were written to individual files. Genes were ranked by AUC to identify top discriminators. To quantify classifier strength across classes, all ROC output files were re-imported and merged into a single data frame containing gene name, associated cell class, and AUC value. Significant classifier genes were defined as those with AUC > 0.75, consistent with prior single-cell ROC classification thresholds.

We next intersected these classifier gene sets with independently generated bulk RNA-seq differentially expressed genes (DEGs). Using DEGs from two comparisons (T26 vs T24 and T22 vs T24), we retained genes with |$\log_2$ fold change ($\log_2$FC)| ≥ 0.33 and adjusted $p < 0.05$ and mapped their fold-changes onto ROC-derived classifier genes. Class-specific gene counts (AUC > 0.75) were computed per comparison, and the distribution of mapped DEGs was visualised using bar plots, jitter plots, and AUC histograms. Chi-square tests were used to compare the frequency of classifier-gene mapping across cell classes. This integrated analysis allowed quantification of which retinal classes contributed most strongly to observed bulk transcriptomic signatures.

Module score projection from bulk DEGs to MRCA

To evaluate how bulk RNA-seq signatures distribute across retinal cell classes at single-cell resolution, we projected up-regulated and down-regulated DEG sets from each bulk comparison onto the MRCA using Seurat's AddModuleScore. For each comparison, two gene modules were defined: (1) up-regulated genes ($\log_2FC \geq 0.33$ & adjusted $p < 0.05$) and (2) down-regulated genes ($\log_2FC \leq -0.33$ & adjusted $p < 0.05$). Module scores were computed with nbin = 10, generating two module-score vectors per dataset. Module score matrices were extracted and standardised (z-scored) across all cells. Scores were then aggregated by major retinal class and summarised as mean ± 1 SD. We visualised class-level projection patterns using DotPlots, line-range plots, and heatmap-style displays, which highlighted how strongly each cell class reflected the “up” and “down” bulk transcriptional programs.

This projection enabled direct interpretation of bulk RNA-seq results in terms of underlying cellular contributors, identifying the cell classes most associated with each transcriptional shift. Comparisons were visualised using faceted dot plots and z-score line-range summaries, allowing side-by-side evaluation of “up” and “down” signatures.

<u>Statistical analysis of data derived from animal experiments</u>

All data from animal experiments, except transcriptome profiling, were analysed using GraphPad Prism (version 10). The animal was the unit of analysis, and the parameters from both eyes were averaged. Data are presented as mean ± SEM, as specified in the figure legends. Comparisons among three or more groups for a single variable were performed using one-way analysis of variance (ANOVA) with Dunnett’s correction for multiple comparisons. For experiments involving two independent variables, two-way ANOVA with Šidák’s correction for multiple comparisons was applied. Longitudinal data with repeated measurements were analysed using repeated-measures ANOVA; when missing values were present, a mixed-effects model (restricted maximum likelihood), as implemented in Prism, was used. The specific statistical tests applied are indicated in the corresponding figure legends. A $p$ value < 0.05 was considered statistically significant.

### Supplementary Figures

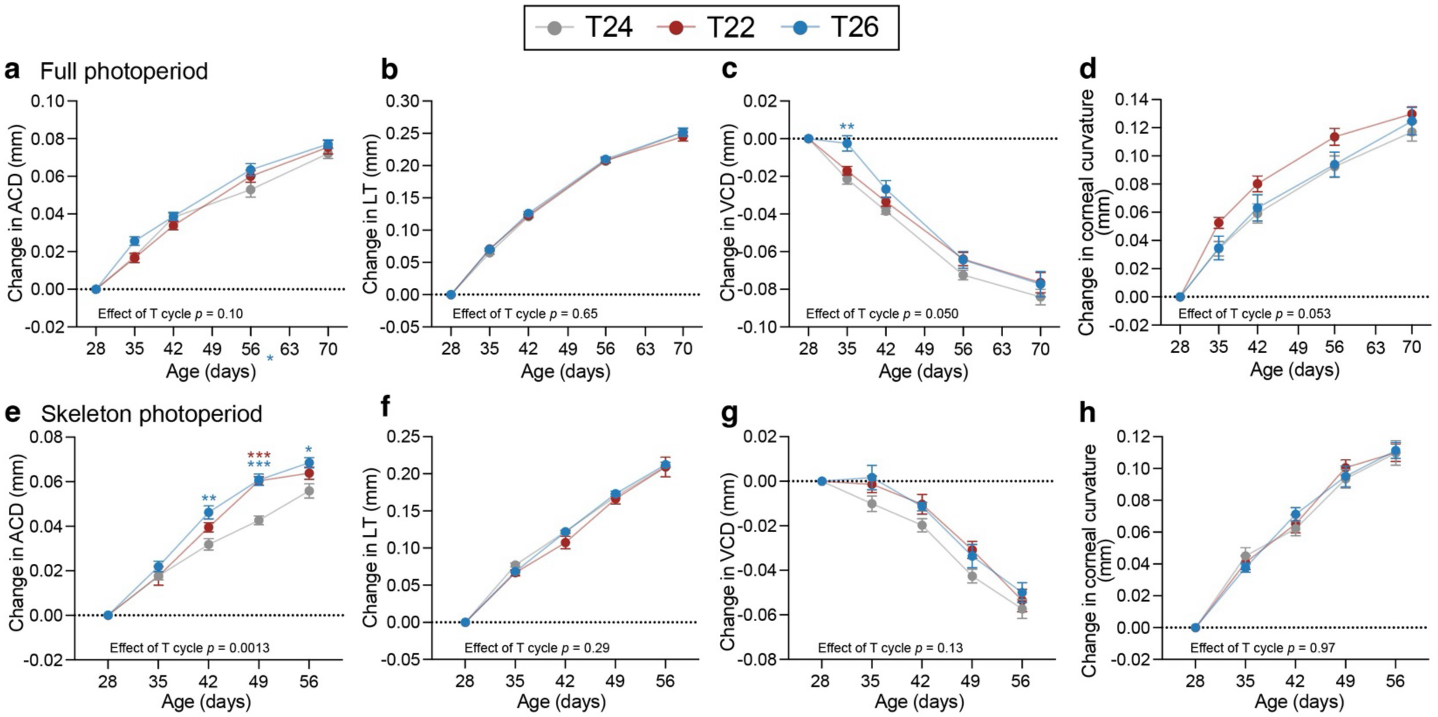


**Supplementary Figure 1. Ocular biometry and corneal curvature under full and skeleton photoperiods of T22 and T26.**

(a-d) Longitudinal changes from baseline (P28) in anterior chamber depth (ACD) (a), lens thickness (LT) (b), vitreous chamber depth (VCD) (c), and corneal curvature (d) in mice housed under T24, T22, and T26 full photoperiods. (e-h) The same measurements under the corresponding skeleton photoperiods, in which 1-h light pulses marked subjective dawn and dusk while maintaining T cycle length. The dashed line indicates no change from baseline. Data are shown as mean ± SEM (full photoperiod: n = 13-14; skeleton photoperiod: n = 6). Statistical analyses were performed using mixed-effects models with Dunnett's correction for multiple comparisons, testing T22 and T26 against T24 controls; *p* values shown in each panel indicate the main effect of T cycle length. * $p < 0.05$, ** $p < 0.01$, *** $p < 0.001$.

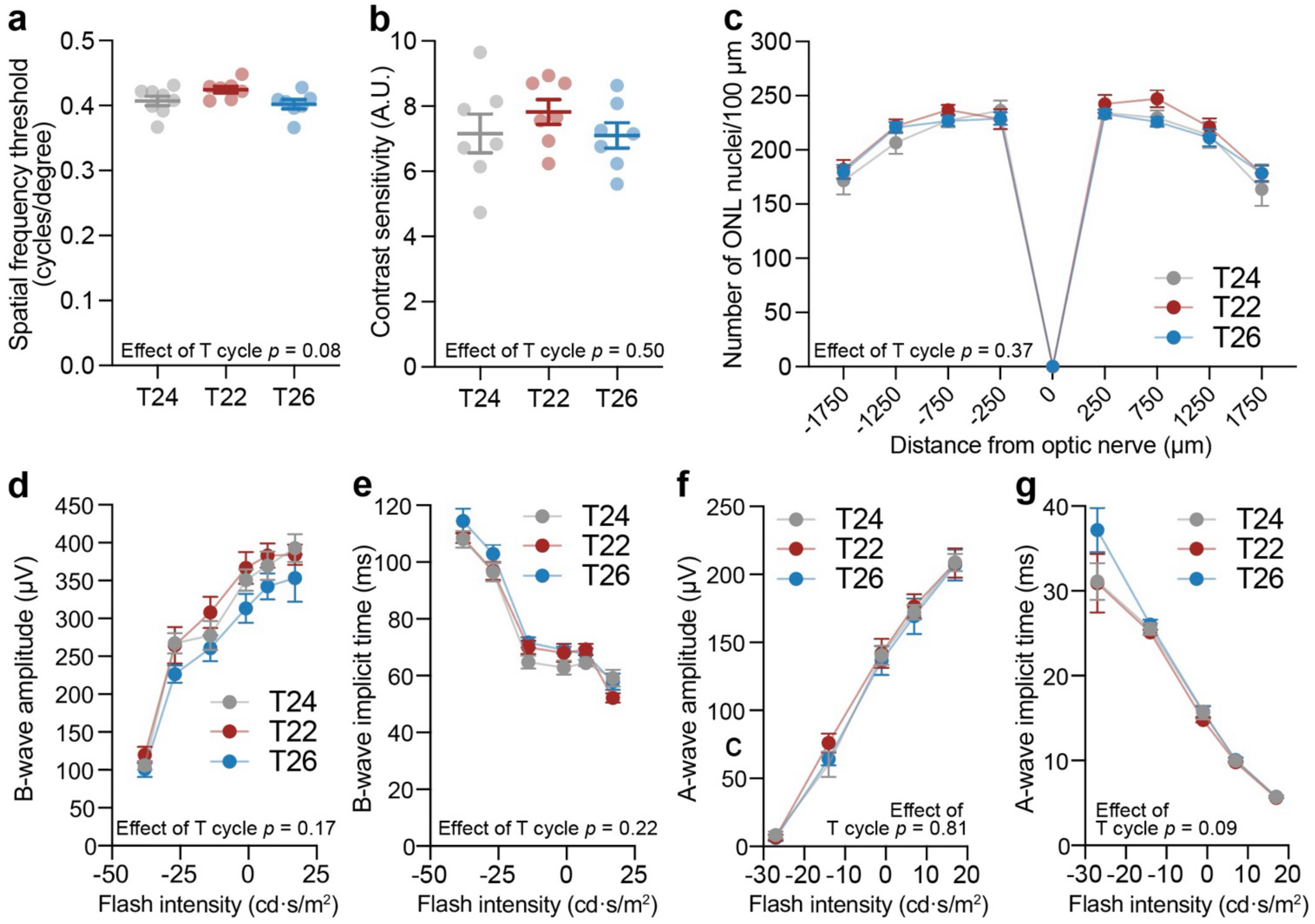


**Supplementary Figure 2. Visual and retinal function in mice housed in T22 and T26.** (a) Visual acuity, measured as the maximum spatial frequency (cycles per degree) eliciting an optokinetic tracking response, in mice housed under T24, T22, or T26 light-dark cycles (n = 7 per group). (b) Contrast sensitivity, measured at 0.103 cycles per degree in the same groups (n = 7 per group). (c) Quantification of outer nuclear layer (ONL) nuclei from retinal sections of mice housed under T24, T22, or T26 conditions (n = 6-11 per group). (d-g) Dark-adapted electroretinography (ERG) measurements showing (d) b-wave amplitude, (e) b-wave implicit time, (f) a-wave amplitude, and (g) a-wave implicit time as a function of flash intensity for each T cycle condition (n = 7 per group). Bars indicate mean ± SEM. Statistical analyses were performed using one-way ANOVA for panels (a-b) and mixed-effects models for panels (c-g).

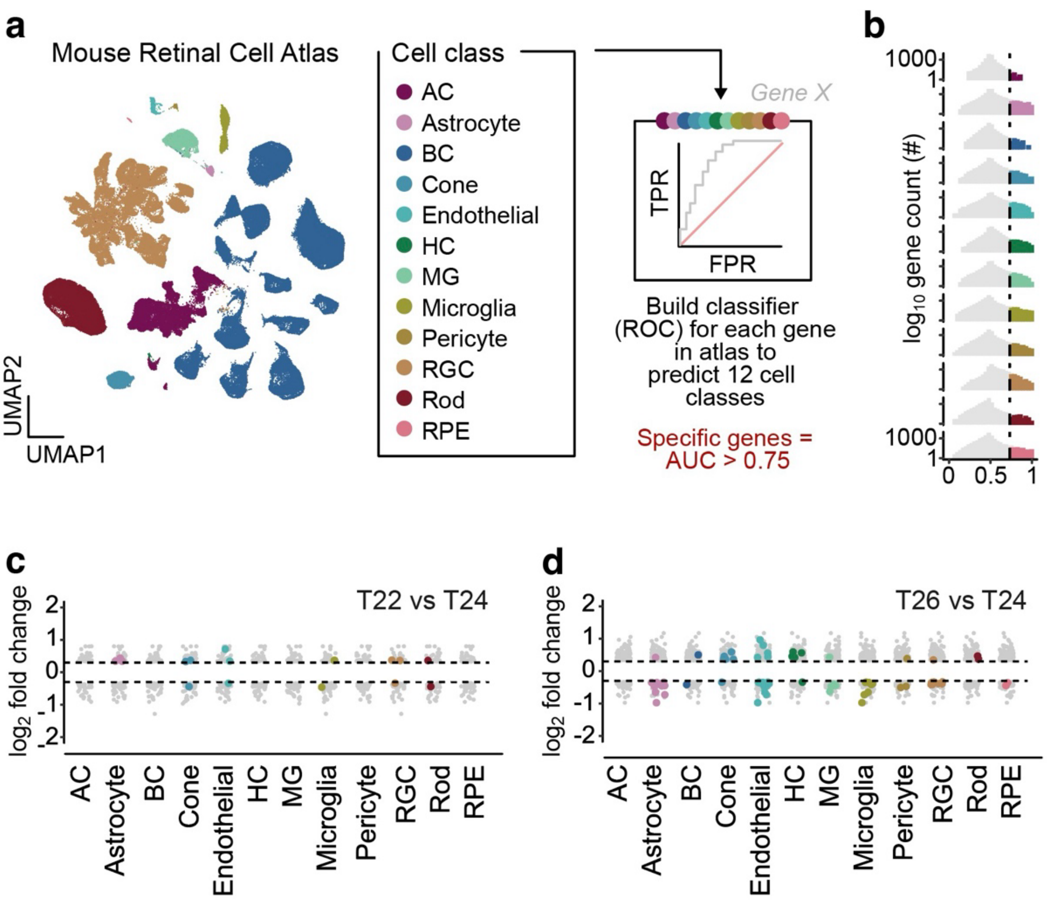


**Supplementary Figure 3. Integration of bulk RNA-seq differential expression with the Mouse Retinal Cell Atlas.**

(a) UMAP embedding of the Mouse Retinal Cell Atlas showing the major retinal neuronal and non-neuronal classes. Each point represents an individual cell; colours correspond to annotated classes. The schematic on the right illustrates the derivation of ROC-based gene-level classifiers for each cell class. (b) Distribution of classifier gene counts across AUC values for all 12 retinal classes. Violin plots show log10 gene counts; coloured regions represent genes with high classifier strength (AUC > 0.75). (c-d) Mapping of bulk DEGs onto classifier genes for the T22 vs T24 (top) and T26 vs T24 (bottom) comparisons. Grey points denote all DEGs; coloured points highlight DEGs that map to high-specificity classifier genes (AUC > 0.75). Dashed lines mark ± 0.33 $\log_2$ fold-change threshold.

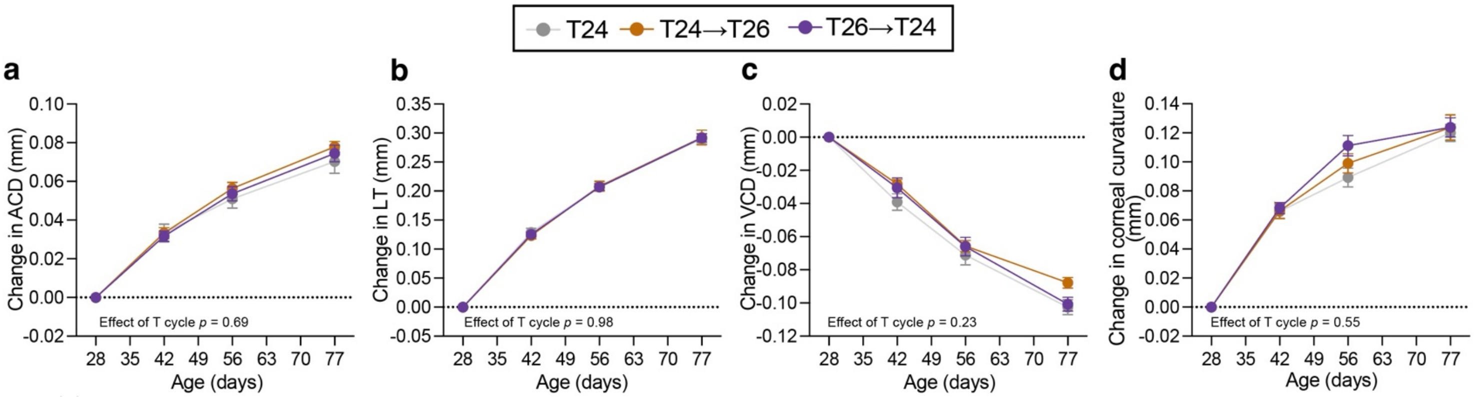


**Supplementary Figure 4. Effects of T26 on ocular biometry and corneal curvature in young adult mice.**

(a-d) Longitudinal changes in anterior chamber depth (ACD) (a), lens thickness (LT) (b), vitreous chamber depth (VCD) (c), and corneal curvature (d) in mice maintained under a standard 24-h light-dark cycle (T24), mice switched from T24 to T26 at P56 (T24→T26), and mice switched from T26 to T24 at P56 (T26→T24). The dashed line indicates no change from baseline. Data are shown as mean ± SEM (n = 11-12 per group). Statistical analyses were performed using mixed-effects models with Dunnett's correction for multiple comparisons, testing T22 and T26 against T24 controls; *p* values shown in each panel indicate the main effect of T cycle length.

### Supplementary Tables

**Supplementary Table 1. Estonian Biobank cohort filtering steps**

**Initial sample size: 137,022 completed questionnaires**

| Exclusion criterion | Resulting sample size (N) |
|---|---|
| Non-standard shift work | 119,373 |
| Waketime < bedtime | 117,033 |
| Bedtime >8h and <15h | 117,033 |
| Waketime >18h and <24h | 117,033 |
| Midsleep time <23 and >34 | 116,532 |
| Sleep latency >150 min | 116,310 |
| Sleep duration <3 or >14h | 116,018 |
| Multiple questionnaires per participant: only the first questionnaire was retained | 114,141 |
| No healthcare data available | 114,067 |
| Concurrent diagnoses of myopia and hyperopia | 110,773 |
| Age >80 | **109,461** |
